# Evaluation of tuberculosis treatment response with serial C-reactive protein measurements

**DOI:** 10.1101/400572

**Authors:** Douglas Wilson, Mahomed-Yunus S Moosa, Ted Cohen, Patrick Cudahy, Collen Aldous, Gary Maartens

## Abstract

**Background:** Novel biomarkers are needed to assess response to antituberculosis therapy in smear-negative patients.

**Methods:** To evaluate the utility of CRP in monitoring response to antituberculosis therapy we conducted a post-hoc analysis on a cohort of adults with symptoms of tuberculosis and negative sputum smears in a high tuberculosis and HIV prevalence setting in KwaZulu-Natal, South Africa. Serial changes in CRP, weight, and haemoglobin were evaluated over 8 weeks.

**Results:** 421 participants with suspected smear-negative tuberculosis were enrolled and 33 excluded. 295 were treated for tuberculosis (137 confirmed, 158 possible), and 93 did not have tuberculosis. 185 of 215 (86%) participants who agreed to HIV testing were HIV-positive. At week 8, the on-treatment median CRP reduction in the tuberculosis group was 79.5% (IQR 25.4; 91.7), median weight gain 2.3% (IQR −1.0; 5.6), and median haemoglobin increase 7.0% (IQR 0.8; 18.9) (p-value <0.0001 for baseline to week 8 comparison of absolute median values). Only CRP changed significantly at week 2 (median reduction of 75.1% (IQR 46.9; 89.2) in the group with confirmed tuberculosis and 49.0% (IQR −0.4; 80.9) in the possible tuberculosis group. Failure of CRP to reduce to ≤55% of the baseline value at week 2 predicted hospitalization or death in both tuberculosis groups, with 99% negative predictive value.

**Conclusion:** Change in CRP may have utility in early evaluation of response to antituberculosis treatment and to identify those at increased risk of adverse outcomes.

**Key points:** C-reactive protein (CRP) falls by 80% after eight weeks of antituberculosis treatment. At two weeks sustained CRP elevation is associated with death or hospitalization.

## Background

Tuberculosis remains a leading cause of death in high HIV prevalence settings, notwithstanding expanding access to antiretroviral therapy (ART), and improved screening strategies and cure rates [1,2]. Widespread implementation of a nucleic acid amplification assay, the Xpert^®^MTB/RIF (GXP), and more recently the Xpert^®^MTB/RIF Ultra, has enhanced rapid laboratory diagnosis of tuberculosis [3,4]. Neither version of the GXP assay however can be used as a proxy measure of treatment response due to persistence of mycobacterial DNA after on-treatment culture conversion [5–7]. Empiric antituberculosis therapy in GXP-negative HIV-positive adults has been frequent, based on non-specific clinical findings, and is likely to persist [8,9]. Current WHO guidelines recommend that all patients are monitored to assess response to therapy, including persistence or reappearance of symptoms of tuberculosis [9]. Sputum smear for acid-fast bacilli (AFB) remains the objective response to therapy criterion, even in the GXP era [5]. However, for patients who are sputum smear-negative at baseline, which occurs frequently in HIV-positive adults, WHO recommend only clinical monitoring, stating “body weight is a useful progress indicator” without specifying criteria [10]. Case fatality rates during antituberculosis treatment are about six-fold higher in HIV co-infected patients, and at least two-fold higher in smear-negative cases [11,12]. Clinicians therefore need specific tools to monitor smear-negative treatment response to detect patients who are not objectively responding to antituberculosis therapy [13].

Several immune-based biomarkers have been proposed to monitor tuberculosis treatment response, including C-reactive protein (CRP), which is an acute inflammatory protein and component of the innate immune response [14–19]. Hepatic synthesis of CRP is induced by interleukin-6 and interleukin-1β [20,21]. Serum or whole blood CRP concentrations are widely used in routine clinical practice and can be measured using affordable point-of-care (POC) devices. Utility of both laboratory and POC CRP testing has been shown when screening tuberculosis suspects in Africa [22–25], with elevated levels in >90% of HIV infected individuals at time of tuberculosis diagnosis [25]. Reductions in CRP concentration after two months on antituberculosis therapy have been described in tuberculosis patients in South Africa, the Gambia, and Brazil [18,26–28]. Recently, a study of 20 HIV positive adults with multi-drug resistant tuberculosis found that sustained on-treatment CRP elevation was associated with increased risk of death [29].

To further evaluate the potential prognostic value of serial CRP measurements we conducted a post-hoc analysis of a prospective cohort of individuals with symptoms of tuberculosis with negative sputum smears or inability to produce sputum, in KwaZulu-Natal, South Africa.

## Methods

### Study population

We conducted a prospective cohort study between June 2005 and February 2007 of adults (over the age of 18 years), with symptoms suggesting tuberculosis and negative sputum smears for AFB, or inability to produce sputum. Potential participants were referred to the study team by healthcare providers at KwaZulu-Natal uMgungundlovu District facilities in the Edendale Hospital catchment area. HIV prevalence in this setting is 16.9%, tuberculosis incidence 922 cases per 100,000, and the rate of HIV coinfection in patients with tuberculosis is around 70% [30–32]. Consenting adults with three sputum smears negative for AFB or without sputum were included in the study if, at the screening visit, constitutional symptoms or cough were reported as being present for >2 weeks or a focal disease process compatible with active tuberculosis was detected. Exclusion criteria were pregnancy, Karnofsky performance score <40, suspected tuberculous meningitis, ≥1 week of antituberculosis treatment, <3 months of antiretroviral therapy (ART), or use of a fluoroquinolone within the past 8 weeks. Additional study details can be obtained from published reports [22,33].

At baseline, study clinicians used standardized criteria with clinical evaluation, chest radiography, and, where indicated, abdominal and pericardial ultrasound scan, to diagnose smear-negative tuberculosis, with initiation of empiric antituberculosis therapy (treatment arm), or to place the participants under observation without antituberculosis therapy (observation arm). The decision whether or not to initiate antituberculosis therapy was made at the baseline visit, before CRP results were available. Sputum and urine specimens for mycobacterial culture were obtained at the baseline visit. Participants’ symptoms were recorded, and functional status estimated using the Karnofsky Performance Score (KPS). Weight was measured in kilograms on a calibrated electronic scale, and C-reactive protein (CRP) and haemoglobin were measured in venous blood at a commercial laboratory.

### On-study clinical review

Standardized clinical review was conducted in all participants at 2, 4 and 8 weeks after the baseline visit, which included KPS, measurement of weight, CRP, and haemoglobin concentration. A Symptom Score Ratio (SSR) was calculated by dividing the aggregate number of symptoms reported as either markedly improved or resolved by the total number of symptoms reported at baseline. Participants who deteriorated clinically were referred for further evaluation by the Edendale Hospital Internal Medicine service. Participants in the observation arm were started on antituberculosis therapy after a pre-treatment baseline evaluation if either a positive mycobacterial culture result became available or if clinical deterioration occurred. These participants (re-baselined group) were then followed up on treatment for a further 8 weeks and on-treatment data were included in the treatment group.

Participants who missed a scheduled appointment were contacted telephonically and rescheduled; these participants’ data were included in the analysis if their visit was within 7 days of their scheduled visit for week 2, and 14 days for weeks 4 and 8. Participants were offered HIV testing at each visit and those with a positive test were referred for antiretroviral therapy at the week 8 visit in keeping with national policy at the time of the study.

At the end of the 8-week follow up period participants were classified as either having confirmed tuberculosis (defined as a positive culture for *M. tuberculosis* complex from any site, or AFB with granulomata on histology); possible tuberculosis (defined as a clinical decision to initiate antituberculosis therapy, without a positive culture), and as not having tuberculosis (defined as negative mycobacterial cultures and no initiation of antituberculosis therapy).

An adverse clinical event was defined as death or hospitalization for a medical condition.

### Statistical analysis

Data were exported from the original Microsoft Access database and analysed using Analyse-it for Microsoft Excel Version 4.51. Distribution of continuous data was determined using the Shapiro-Wilk test. Groups of unpaired non-parametric data were compared using the Wilcoxon-Mann-Whitney test. Changes in potential response to therapy measures over time were compared using the Friedman test for repeated measurements; participants with missing data points were excluded from the model. Data obtained from participants classified with an adverse outcome after week 2 were retained in the analysis. Change in CRP was calculated by subtracting the week 2 or week 8 value from the baseline value, and percentage change by dividing the change by the baseline value. Change in haemoglobin concentration or weight was calculated by subtracting the baseline measurement from the week 2 or week 8 measurement and dividing the change by the baseline value. Sensitivity / specificity decision tables were generated and used to select CRP percentage change cut-offs with a sensitivity of >90% for death or hospitalization. Binary nominal variables were compared using Fisher’s exact test, and 2×2 tables to calculate test performance characteristics. Confidence intervals were set at 95%; those for area under the receiver operating characteristic curve were calculated using R, version 3.4.1, Vienna, Austria.

### Ethics review

The study was approved by ethics review boards of the Universities of KwaZulu-Natal and Cape Town, and by the KwaZulu-Natal Department of Health. All participants provided written informed consent.

## Results

Four hundred and twenty-one participants were enrolled and 388 (92.6%) were included in this analysis. Baseline characteristics are shown in in Table 1. Five participants (1.3%) were taking antiretroviral therapy at the baseline visit. Sixteen participants initially included in the observation arm were diagnosed with tuberculosis and were started on antituberculosis treatment: 13 with confirmed tuberculosis, and 3 diagnosed empirically. Participant exclusions and outcomes over the 8-week follow-up period are shown in Fig. 1: 295 (76.0%) participants started on antituberculosis therapy during the study period and 93 (24.0%) were observed.

**Table 1:** Baseline characteristics of the 388 participants evaluated for tuberculosis

| Characteristic |  |
| --- | --- |
| Age median (IQR) years | 34.4 (29.5; 41.8) |
| Male n (%) | 212 (55.9) |
| HIV positive n (%) | 185 (47.4) |
| HIV negative n (%) | 30 (7.7) |
| Declined HIV testing n (%) | 175 (44.9) |
| Received antibiotic within 2 weeks before enrolment n (%) | 280 (71.8) |
| Cough for >2 weeks n (%) | 363 (93.1) |
| Weight loss n (%) | 295 (75.6) |
| Anorexia n (%) | 275 (70.5) |
| Night sweats n (%) | 263 (67.4) |
| Fatigue n (%) | 246 (63.1) |
| Fever and chills n (%) | 227 (58.2) |
| Chest pain n (%) | 221 (56.7) |
| Haemoptysis n (%) | 44 (11.3) |

**Figure 1.**
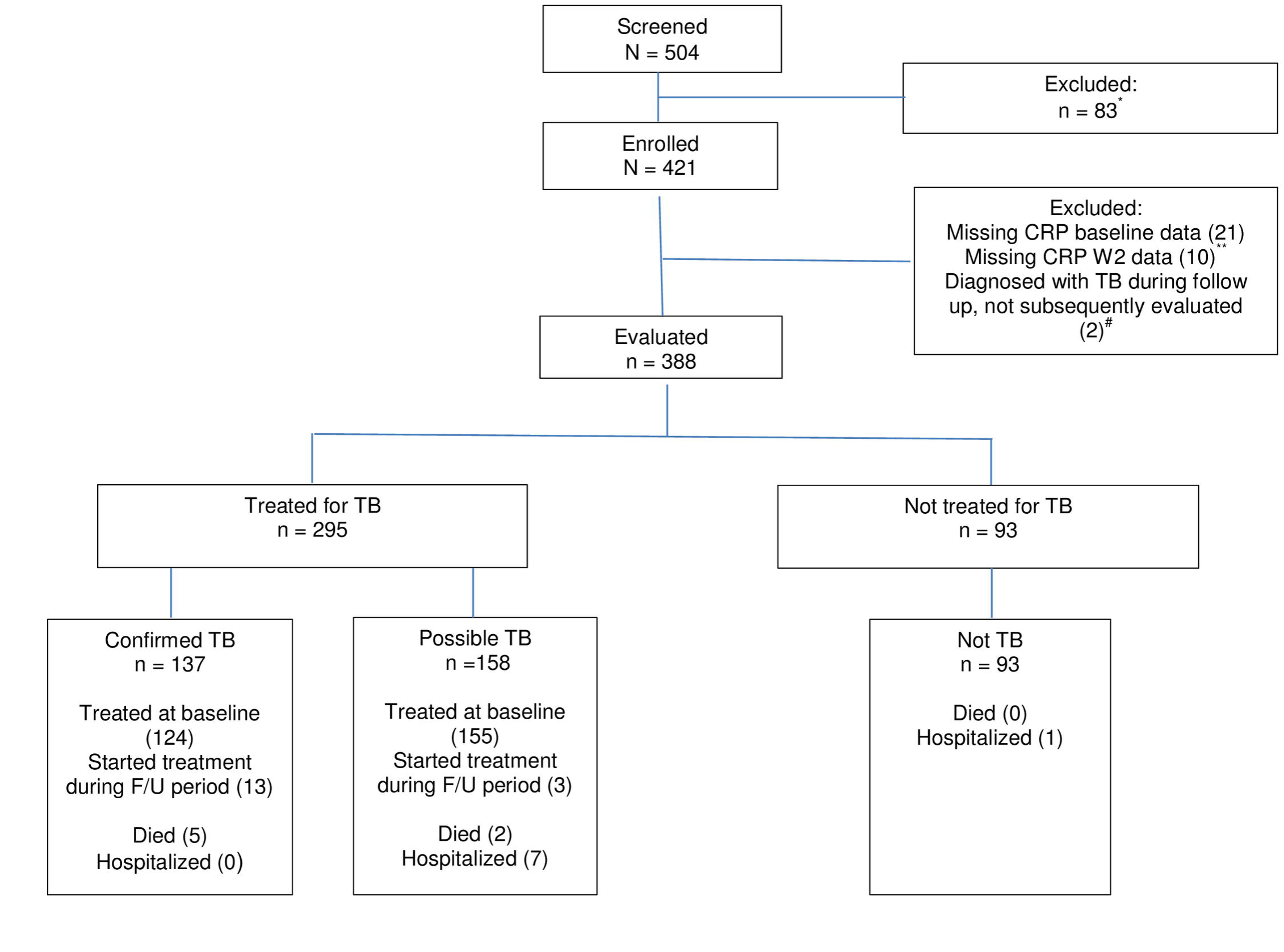
Participant outcomes during the 8-week follow up period. *Not able to attend for regular review (28), no active symptoms (17), alternative medical diagnosis (14), Karnofsky score <40 (5), pneumocystis pneumonia (4), informed consent not obtained (3), sputum smear-positive (3), already on anti-tuberculosis treatment (3), other (6). **Confirmed TB (4); possible TB (2); not TB (3); TB diagnosed on culture not treated at BL visit (1). #Died before antituberculosis treatment initiation (1); referred for inpatient Rx before re-baselined week 2 visit (1). TB = tuberculosis.

### Changes in response to therapy parameters during the eight-weeks follow up period

Table 2 shows trends for CRP, weight, haemoglobin, KPS and SSR stratified by final diagnosis. CRP showed the greatest changes from baseline and the greatest early change in the on-treatment tuberculosis groups. Only the objectively measured response to therapy parameters (CRP, weight, and haemoglobin) showed no significant change over the 8 week follow up period in the group not treated for tuberculosis.

**Table 2:** Trends in clinical parameters in participants treated for tuberculosis, with confirmed tuberculosis, possible tuberculosis, and not treated for tuberculosis. Values at different timepoints are medians with interquartile ranges in parentheses.

|  |  | n* | Baseline | Week 2 | Week 4 | Week 8 | p-value<sup>#</sup> |
| --- | --- | --- | --- | --- | --- | --- | --- |
| C-Reactive Protein mg/L | Treated for TB | 265 | 56.2 (17.6; 114.2) | 13.0 (4.0; 33.3) | 14.0 (5.0; 28.4) | 8.6 (3.0; 20.0) | <0.0001 |
|  | Confirmed TB | 125 | 84.0 (45.8; 128.3) | 15.8 (7.2; 39.5) | 18.0 (7.0; 35.7) | 8.0 (3.0; 20.2) | <0.0001 |
|  | Possible TB | 140 | 30.4 (8.0; 84.1) | 10.8 (3.0; 26.8) | 9.6 (4.0; 24.0) | 9.0 (3.0; 20.0) | <0.0001 |
|  | Not treated for TB | 82 | 3.9 (2.0; 10.0) | 3.0 (2.0; 9.0) | 3.0 (2.0; 10.1) | 3.4 (2.0; 9.0) | 0.5 |
| Weight kg | Treated for TB | 270 | 56.8 (50.1; 64.4) | 57.0 (50.6; 64.9) | 57.9 (51.2; 65.5) | 58.4 (52.3; 65.8) | <0.0001 |
|  | Confirmed TB | 128 | 55.6 (49.7; 62.1) | 55.5 (50.2; 62.1) | 56.6 (50.6; 62.1) | 57.7 (52.5; 63.5) | <0.0001 |
|  | Possible TB | 142 | 58.1 (50.5; 66.0) | 58.9 (51.0; 66.6) | 59.0 (51.9; 66.8) | 59.9 (52.2; 68.1) | <0.0001 |
|  | Not treated for TB | 82 | 59.9 (53.1; 67.6) | 60.4 (53.9; 68.2) | 61.0 (54.2; 69.3) | 60.4 (54.6; 69.8) | 0.4 |
| Haemoglobin g/dL | Treated for TB | 265 | 10.6 (9.2; 12.1) | 10.6 (9.4; 12.2) | 11.1 (9.8; 12.4) | 11.7 (10.2; 12.9) | <0.0001 |
|  | Confirmed TB | 125 | 10.2 (8.8; 11.7) | 10.4 (9.2; 11.7) | 10.8 (9.7; 12.3) | 11.7 (10.3; 12.8) | <0.0001 |
|  | Possible TB | 140 | 11.1 (9.7; 12.4) | 10.9 (9.6; 12.4) | 11.2 (10.0; 12.7) | 11.7 (10.0; 13.0) | <0.0001 |
|  | Not treated for TB | 80 | 13.3 (11.4; 14.4) | 13.0 (11.6; 14.4) | 13.0 (11.3; 14.3) | 13.2 (11.7; 14.5) | 0.4 |
| Karnofsky Performance Score | Treated for TB | 270 | 70 (60; 80) | 70 (70; 80) | 80 (70; 80) | 90 (80; 90) | <0.0001 |
|  | Confirmed TB | 128 | 60 (50; 70) | 70 (60; 80) | 80 (70; 80) | 80 (80; 90) | <0.0001 |
|  | Possible TB | 142 | 70 (60; 80) | 80 (70; 80) | 80 (80; 90) | 90 (80; 90) | <0.0001 |
|  | Not treated for TB | 82 | 70 (70; 80) | 80 (70; 80) | 80 (70; 80) | 80 (80; 90) | <0.0001 |
| Symptom Score Ratio | Treated for TB | 270 | 0.0 (0.0; 0.0) | 0.8 (0.6; 1.0) | 1.0 (0.8; 1.0) | 1.0 (1.0; 1.0) | <0.0001 |
|  | Confirmed TB | 128 | 0.0 (0.0; 0.0) | 0.8 (0.6; 1.0) | 0.9 (0.8; 1.0) | 1.0 (0.9; 1.0) | <0.0001 |
|  | Possible TB | 142 | 0.0 (0.0; 0.0) | 0.8 (0.7; 1.0) | 1.0 (0.8; 1.0) | 1.0 (1.0; 1.0) | <0.0001 |
|  | Not treated for TB | 82 | 0.0 (0.0; 0.0) | 0.6 (0.3; 0.8) | 0.8 (0.5; 1.0) | 1.0 (0.7; 1.0) | <0.0001 |

\*Number of participants contributing complete data set to model. <sup>#</sup>p-value for trend

At week 2, median percentage CRP reduction was 62.7% (IQR 19.7; 85.1) in the entire group treated for tuberculosis: 75.1% (IQR 46.9; 89.2) in the confirmed tuberculosis group and 49.0% (IQR −0.4; 80.9) in the possible tuberculosis group. By week 8, median percentage CRP reduction was 79.5% (IQR 25.4; 91.7) in the group treated for tuberculosis, 86.3% (IQR 74.8; 92.4) in the confirmed tuberculosis group, 65.9% (IQR −26.6; 86.7) in the possible tuberculosis group.

Median percentage CRP reduction was similar in participants with known HIV status treated for tuberculosis: HIV positive vs. HIV negative 58.1% (IQR 3.9; 84.3) vs. 23.8% (IQR - 40.4; 90.9) at week 2 (p-value 0.3), and 79.9% (IQR 13.7; 91.9) vs. 44.4% (IQR −19.6; 89.2) at week 8 (p-value 0.3).

In the group not treated for tuberculosis, the median CRP change did not change significantly: 0.0% (IQR −36.2; 24.5) at week 2, and 0.0% (IQR −26.0; 63.2) at week 8.

In the group that did not start antituberculosis therapy at baseline, the median CRP in the sub-group subsequently diagnosed with confirmed tuberculosis was similar to the sub-group not diagnosed with tuberculosis (8.9 mg/L (IQR 2.0; 42.8) vs. 3.0 mg/L (IQR 2.0; 8.9) p-value 0.2); this similarity persisted through Week 4 (3.0 mg/L (IQR 3.0; 6.2) vs. 3.0 mg/L (2.0; 10.8) p-value 1.0).

Weight and haemoglobin improved significantly (p-value <0.0001) by week 8 in the confirmed and possible tuberculosis groups, but the magnitude of change was modest, and changes at week 2 were not significant. At week 8, weight increased by 2.3% (IQR −1.0; 5.6) in the entire group treated for tuberculosis, by 3.6% (IQR −0.7; 7.2) in the group with confirmed tuberculosis; 1.6% (IQR −1.4; 4.3) in the group with possible tuberculosis; and 0.7% (IQR −1.5; 2.9) in the group without tuberculosis. Haemoglobin increased by 7.0% (IQR 0.8; 18.9) in the entire group treated for tuberculosis, 10.0% (IQR 3.0; 22.9) in the group with confirmed tuberculosis; 3.6% (IQR −1.8; 15.4) in the group with possible tuberculosis; and 0.0% (−3.2; 5.9) in the group without tuberculosis.

The subjectively assessed KPS and SSR parameters improved both in the tuberculosis group, and in the group not treated for tuberculosis. In comparison to the tuberculosis group however, the week 8 KPS improved less in the group without tuberculosis, and improvement in the SSR occurred at a slower rate and was less complete. To further evaluate these observations the group treated for tuberculosis and with paired data (n = 272) was compared with the group without tuberculosis and paired data (n = 82). The week 8 KPS improved by a median of 20 (IQR 10; 30) in the tuberculosis group and by 10 (IQR 0; 10) in the group without tuberculosis; the week 8 SSR reached a median of 1.0 (IQR 1.0; 1.0) in the tuberculosis group and 1.0 (IQR 0.7; 1.0) in the group without tuberculosis (p-value <0.0001 for both comparisons).

### Change of response to therapy parameters from baseline to week 2 as predictors of adverse outcomes in the tuberculosis group

Adverse outcomes in the group treated for tuberculosis were defined as either death (n=7) or hospitalization (n=7) during the study period (Fig. 1). At week 2, both CRP and the SSR showed meaningful improvement in the group treated for tuberculosis, and percentage improvement was associated with death or hospitalization (Table 3). The area under the receiver operating characteristic curve was similar for change in CRP and change in SSR (0.83 [95% IQR 0.70; 0.95] vs. 0.75 [95% IQR 0.60; 0.90], difference 0.08 [95% CI −0.12; 0.27]).

**Table 3:** Median percentage change in RTT parameters from baseline to week 2 and AROC for predicting death or hospitalization in the tuberculosis treatment group. n = 295.

|  | Percentage change from baseline (IQR) | AROC (95% CI) for hospitalization or death |
| --- | --- | --- |
| C-Reactive protein | 62.7 (9.7; 85.1) | 0.83 (0.70; 0.95) |
| Symptom Score Ratio | 80 (60; 100) | 0.75 (0.60; 0.90) |
| Haemoglobin | 1.1 (-3.8; 6.9) | 0.70 (0.55; 0.84) |
| Weight | 0.5 (-1.1; 2.4) | 0.65 (0.49; 0.81) |
| Karnofsky Performance Score | 14.2 (0.0; 16.7) | 0.57 (0.40; 0.75) |

RTT: response to therapy; AROC: area under receiver operating characteristic curve; IQR:
interquartile range; CI: confidence interval

The median CRP reduction from baseline to week 2 was 31.0 mg/L (IQR 3.3; 77.1) in the 281 participants with an uncomplicated clinical outcome (65.1% reduction [IQR 19.9; 85.4]), and −6.0 mg/L (IQR −56.5; 1.0) in the 14 who died or were hospitalized (−33.8% reduction [IQR −198.8; 4.3], (p-value <0.0001 for both comparisons). For the SSR, the median score was 0.8 (IQR 0.63; 1.0) in the participants with an uncomplicated course, and 0.5 (IQR 0.21; 0.62) in those with an adverse outcome (p-value 0.001).

Receiver operating characteristics for the combined endpoint of death or hospitalization vs. percentage week 2 CRP change for participants treated for tuberculosis, and in the confirmed and possible tuberculosis sub-groups, are shown in Fig 2. Table 4 shows the performance characteristics of the percentage change in CRP at week 2, at a cut-off of ≤55%, used as a test to detect those at risk of death or hospitalization. Overall, at this cut-off, in the entire group treated for tuberculosis, sensitivity was 92%, specificity 59%, and negative predictive value of 99%.

**Table 4:** On treatment performance characteristics of baseline to week 2 change in C-reactive protein in predicting death or hospitalization at percentage change cut-off of ≤55%

|  | Treated for TB | Confirmed TB | Possible TB |
| --- | --- | --- | --- |
| No. participants | 295 | 137 | 158 |
| No. died or hospitalized | 14 | 5 | 9 |
| Sensitivity | 0.92 (0.69; 0.99) | 1.0 (0.57; 1.0) | 0.89 (0.57; 0.98) |
| Specificity | 0.59 (0.53; 0.64) | 0.71 (0.62; 0.78) | 0.48 (0.40; 0.56) |
| Youden's index | 0.52 | 0.71 | 0.37 |
| Positive predictive value | 0.10 (0.08; 0.12) | 0.1 (0.09; 0.14) | 0.09 (0.07; 0.12) |
| Negative predictive value | 0.99 (0.96; 1.0) | 1.0 (-; -) | 0.99 (0.92; 1.0) |
| Positive likelihood ratio | 2.2 (1.6; 2.7) | 3.4 (1.8; 4.5) | 1.7 (1.1; 2.1) |
| Negative likelihood ratio | 0.12 (0.02; 0.53) | 0.0 (0.0; 0.62) | 0.23 (0.04; 0.92) |
| Odds Ratio | 18.5 (3.0; 111.8) | $+\infty$ (3.0; $+\infty$) | 7.5 (1.2; 47.2) |

Figures in brackets = 95% confidence intervals; CRP = C-reactive protein, TB = tuberculosis

**Fig 2:**
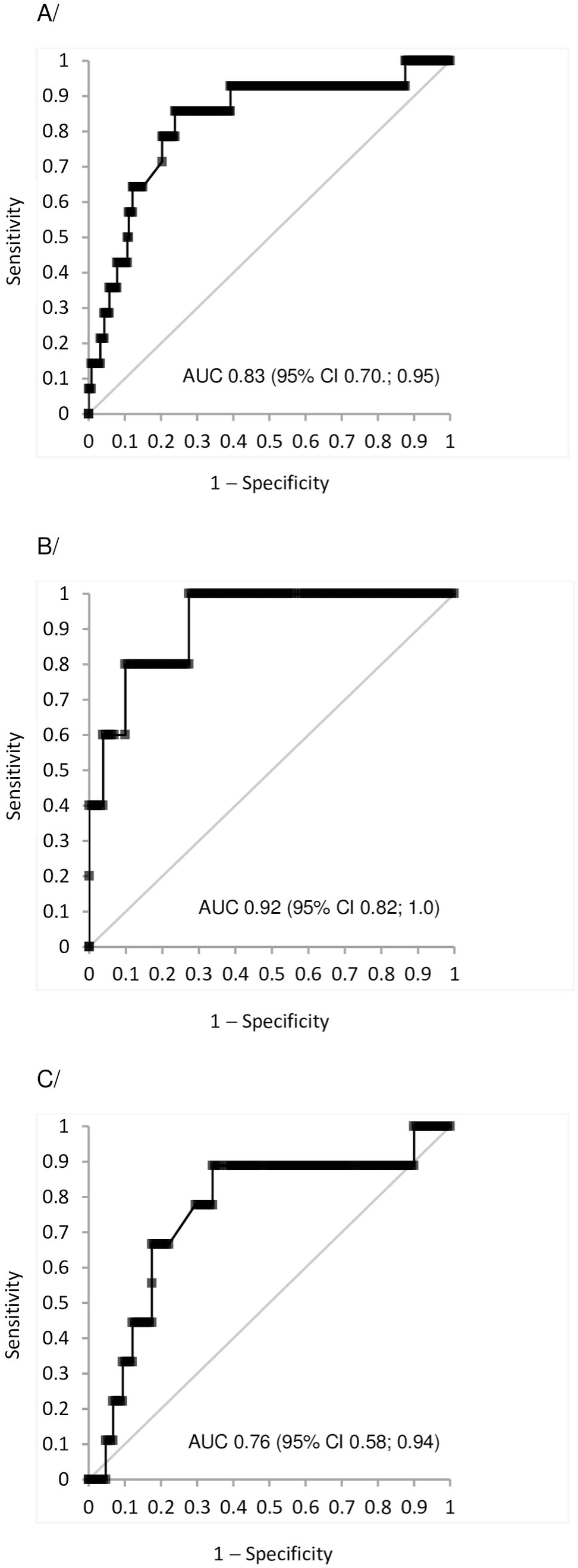
Receiver operating characteristics for percentage change in C reactive protein and death/ hospitalization. A/ Participants treated for tuberculosis (n = 295); B/ Participants with confirmed tuberculosis (n = 137); C/ Participants with possible tuberculosis. (n = 158). AUG area under the receiver operating characteristic curve

## Discussion

Median CRP concentrations followed strikingly different patterns in participants treated for tuberculosis, and in those being observed for tuberculosis. Those in the observation group had low CRP concentrations, which remained unchanged during the study period. In contrast, participants diagnosed empirically with tuberculosis and started on treatment had higher CRP concentrations at baseline, which fell significantly by week 2 and approached the normal range by week 8. The trend was most marked in participants with confirmed tuberculosis. These findings are compatible with other studies [18, 26–29], and add additional insights into the potential for CRP to be used as a tool to evaluate response to antituberculosis therapy.

Change in CRP at week 2 predicted death or hospitalization during 8 weeks of antituberculosis therapy. The group of tuberculosis patients who died or were hospitalized within 8 weeks were less likely to experience a reduction in CRP concentration at week 2 from the baseline concentration. Overall, a decrease in CRP of ≤55% at two weeks predicts death or hospitalization, with a negative predictive value of 99%. However, this finding is imprecise due to the small number of events.

The other objective response to therapy parameters, weight and haemoglobin, have limited utility as they did not change significantly at week 2 and showed only modest improvement at week 8. The subjective response to therapy parameters changed early and with reasonable magnitude, but their utility is limited by significant changes in participants without tuberculosis; they could have value in patients with positive rapid diagnostic tests for tuberculosis. Change in SSR at week 2 had reasonable discrimination in predicting adverse events.

Our study has a number of limitations. First, the proportion of participants with unknown HIV status was high, and number of HIV-infected participants on antiretroviral therapy (ART) at the time of enrolment was low. ART was started only after completion of the 8-week study period in line with guidelines when the study was done. Immune reconstitution syndrome in HIV seropositive patients with tuberculosis at the time of ART initiation is associated with elevated CRP concentrations [34] and may limit the value of change in CRP at week 2 to predict adverse clinical events in patients initiating ART soon after commencing antituberculosis treatment. Second, drug susceptibility testing was not performed in this study, and some participants may have had drug resistant infection not responding to first line treatment. However, in South Africa at the time of this study only about 2% of tuberculosis cases had multidrug resistance [35]. Third, the number of adverse outcomes in those with confirmed tuberculosis was low in comparison to those with possible tuberculosis, suggesting that diagnoses mimicking tuberculosis may have caused adverse outcomes in the tuberculosis treatment group. This would not, however, alter the need for further medical evaluation in this group should CRP remain elevated. Finally, this study was conducted prior to the implementation of GXP as the first line diagnostic test for tuberculosis and CRP trends may be different in GXP-negative tuberculosis cases.

This analysis provides additional information on the utility of monitoring CRP trends to assess early treatment response, with sustained CRP levels at week 2 of treatment being associated with increased risk of adverse clinical outcomes. Further evaluation in the GXP era is needed.

## Acknowledgement

The authors acknowledge the study clinicians Dr Lindo Mbhele and Dr Langa Ngubane, and the research nurses Sr Pat Bartman and Sr Zanele Magcaba. This study was funded by BMS Secure the Future.

